# Accelerated nanopore basecalling with SLOW5 data format

**DOI:** 10.1101/2023.02.06.527365

**Authors:** Hiruna Samarakoon, James M. Ferguson, Hasindu Gamaarachchi, Ira W. Deveson

**Affiliations:** Genomics Pillar, Garvan Institute of Medical Research, Sydney, NSW, Australia; Centre for Population Genomics, Garvan Institute of Medical Research and Murdoch Children’s Research Institute, Australia; School of Computer Science and Engineering, University of New South Wales, Sydney, NSW, Australia; Faculty of Medicine, University of New South Wales, Sydney, NSW, Australia

**Author notes:** Contributed equally (Hiruna Samarakoon, James M. Ferguson). Joint-senior authors; correspondence &.

## Abstract

Nanopore sequencing is emerging as a key pillar in the genomic technology landscape but computational constraints limiting its scalability remain to be overcome. The translation of raw current signal data into DNA or RNA sequence reads, known as ‘basecalling’, is a major friction in any nanopore sequencing workflow. Here, we exploit the advantages of the recently developed signal data format ‘SLOW5’ to streamline and accelerate nanopore basecalling on high-performance computer (HPC) and cloud environments. SLOW5 permits highly efficient sequential data access, eliminating a significant analysis bottleneck. To take advantage of this, we introduce *Buttery-eel*, an open-source wrapper for Oxford Nanopore’s *Guppy* basecaller that enables SLOW5 data access, resulting in performance improvements that are essential for scalable, affordable basecalling.

## INTRODUCTION

Sequencing devices from Oxford Nanopore Technologies (ONT) afford countless opportunities in research and clinical genomics. With the unique ability to sequence both short and long native DNA and RNA molecules on an inexpensive device, ONT presents a significant value proposition, and is already disrupting the genomic technology landscape. To realize large-scale adoption and replacement of dominant short-read sequencing platforms (i.e. Illumina, BGI, IonTorrent), however, bottlenecks in nanopore data analysis must be addressed.

ONT devices measure the displacement of ionic current as a DNA/RNA molecule passes through a nanoscale protein pore, recording time-series signal data that can be translated – or ‘basecalled’ – to determine the molecule’s sequence^1^. Basecalling is the first step in virtually any nanopore analysis workflow. Real-time data acquisition occurs in parallel across thousands of pores on a single ‘flow-cell’ during a sequencing run. This raw data can be basecalled in real-time using the on-board computer on an ONT device, or subsequent to completion of the experiment, using an external computer^1^.

ONT’s high-output PromethION device has enabled increasingly cost-effective nanopore sequencing of human and other large eukaryotic genomes^2–6^, with scalability that theoretically rivals dominant short-read sequencing platforms. The PromethION (P48 version) is equipped to run 48 flow-cells in parallel and has capacity to sequence ~96 human genomes at >30-fold coverage per week (assuming one flow-cell per sample). However, in our experience, the PromethION’s attached compute tower (PRO-PRCV100) is unable to execute live high-accuracy basecalling on more than ~8-10 flow-cells in parallel. Therefore, to realise even a fraction of the machine’s theoretical sequencing throughput, the user must export the raw data and perform basecalling externally on their own high-performance computer (HPC) or cloud environments^3,5,7,8^.

As we have shown previously^9^, ONT’s native data format ‘FAST5’ is large and poorly engineered for efficient analysis on parallel computer systems. These limitations are perhaps most salient during basecalling, where existing inefficiencies result in major costs and impediments for large-scale nanopore projects. For example, as *we* show below, processing a single human genome dataset with ONT’s production basecalling software *Guppy* on a typical Amazon Web Services (AWS) cloud instance takes at least ~13 hours and costs ~$165 USD, or up to ~25% of the cost of the flow-cell used to generate the data.

We recently developed a new file format, SLOW5, which is designed to resolve the inherent limitations in FAST5^9^. In its compressed binary form (BLOW5), the new format is ~20-80% smaller than FAST5 and permits efficient parallel access by multiple CPU threads. We showed previously that SLOW5 enables order-of-magnitude improvements in the speed of common nanopore data analysis processes, such as DNA methylation calling. However, the potential benefits of SLOW5 on basecalling have not been investigated, largely because ONT’s production basecalling software, *Guppy*, does not currently support SLOW5 data access.

Here, we explore the benefits of the SLOW5 format for scalable basecalling of nanopore data. We articulate a new advantage of SLOW5, namely its capacity for rapid sequential data access (as opposed to random access, explored previously^9^), which can be exploited to accelerate basecalling. We also unveil *Buttery-eel*, an open-source wrapper that enables SLOW5 data access by *Guppy*, and demonstrate the resulting performance benefits during nanopore basecalling.

## RESULTS

Basecalling is a major friction in any nanopore sequencing workflow. During basecalling with ONT’s *Guppy* software, raw-signal reads are retrieved from the FAST5 input file, passed to the central processing unit (CPU)s or graphics processing unit (GPU)s, where a neural network model is applied to translate the signal into a DNA sequence, which is then written to a FASTQ output file^10^. File reading is performed in a random access pattern because the complex structure of a FAST5 file does not allow more efficient sequential access. When sufficient CPU/GPU capacity is available, the rate of data access from the FAST5 file, rather than data processing on the CPU/GPU, has the potential to become a bottleneck for the overall analysis.

Binary SLOW5 (BLOW5) format is designed to enable efficient file reading by either random or sequential access patterns, using the *slow5lib* (C++) or *pyslow5* (python) libraries^9^. Raw reads do not need to be basecalled in any particular order, meaning highly efficient sequential access should be preferred. We therefore reasoned that the use of fast, sequential access with BLOW5, instead of slow random access with FAST5, might improve basecalling performance. *Guppy* does not currently support BLOW5 data access and the *Guppy* code cannot be modified directly to enable this (because it is commercial, close-source software). Instead, we developed *Buttery-eel*, a client wrapper to enable basecalling of BLOW5 files with the latest version of *Guppy* (see **Methods**). *Buttery-eel* uses *pyslow5* to access reads sequentially from a BLOW5 file/s, which are then submitted to a *Guppy* server for processing, before writing them to a FASTQ output file (**Figure 1a**). *Buttery-eel* has equivalent usage to *Guppy*, identical outputs, and it maintains all key functionality of the latest *Guppy* version, including DNA methylation calling with *Remora* built-in.

**Figure 1.**
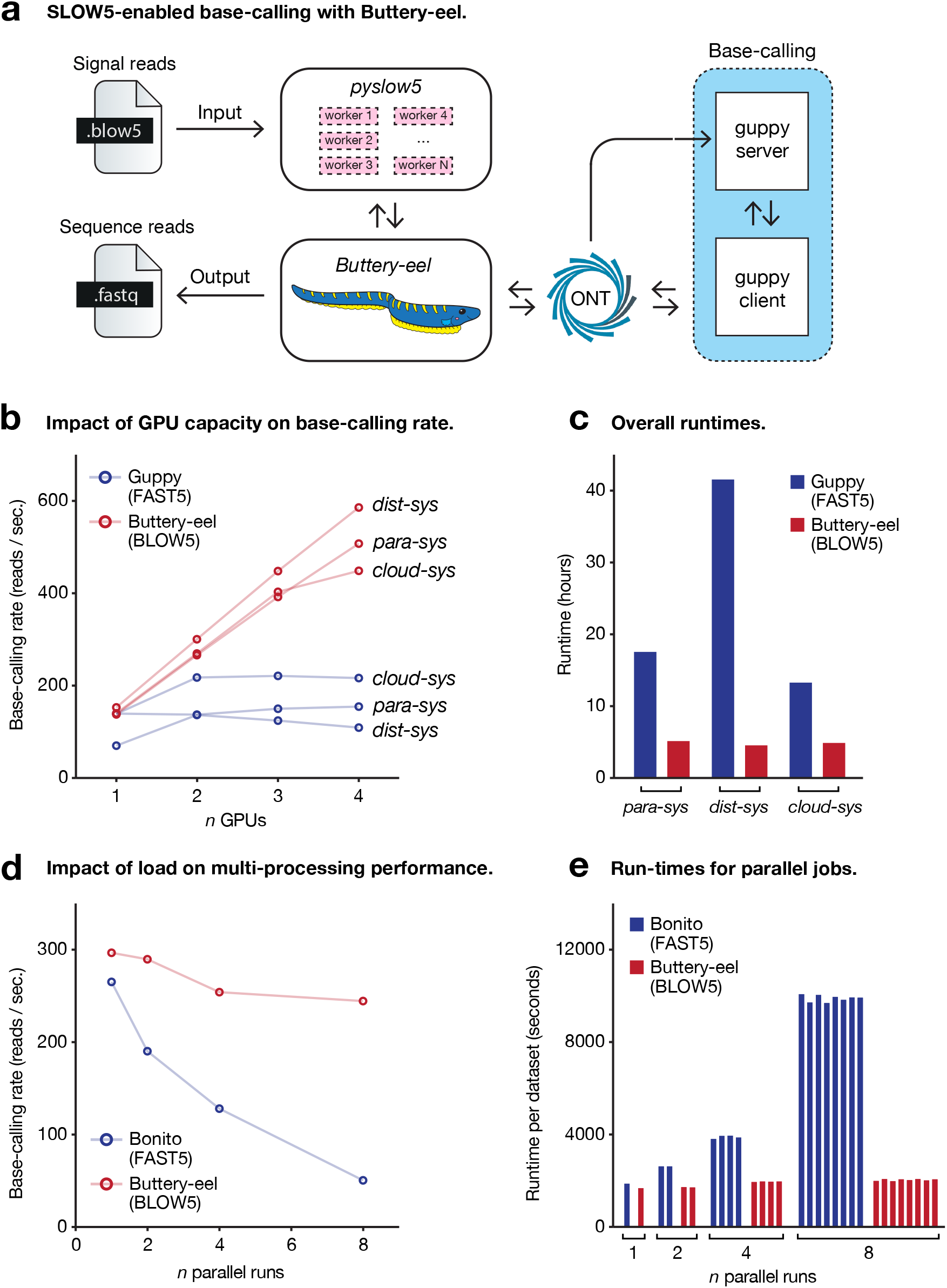
Acceleration of ONT basecalling with Buttery-eel. (**a**) Schematic diagram describing the architecture of *Buttery-eel*, which is a client wrapper that enables input of BLOW5 files to ONT’s *Guppy* basecaller. *Buttery-eel* uses *pyslow5* to access reads from a BLOW5 file. Reads are then submitted to a *Guppy* server for processing, before writing them to a FASTQ (or SAM) output file. (**b**) Line plots show the basecalling rate (in reads/second) achieved by *Guppy* with FAST5 input and Buttery-eel with BLOW5 input, relative to the number of GPUs utilised on each of three computer systems (*dist-sys, para-sys, cloud-sys;* see **Supplementary Table 2**). (**c**) Bar chart shows the minimum execution time achievable (i.e with 4 GPUs) for a typical 30X human genome sequencing dataset processed on each of the systems in **b**. (**d**) Line plots show the basecalling rate achieved by *Bonito* with multiprocessing on FAST5 input and *Buttery-eel* with BLOW5 input, relative to the number of identical basecalling jobs running in parallel. (**e**) Bar chart shows the overall execution time for each individual dataset (500,000 reads), with 1, 2, 4 or 8 basecalling jobs running in parallel.

To evaluate the impact of improved BLOW5 data access on basecalling performance, we processed a small dataset of ~500,000 reads in either FAST5 (with *Guppy*) or BLOW5 (with *Buttery-eel*) format (**Supplementary Table 1**). In both cases we used the identical ‘high-accuracy’ (HAC) basecalling model. The analysis was run on three different multi-GPU HPC/cloud systems, which represent a range of architectures commonly used by the genomics community, specifically: (*i*) an institutional HPC with a parallel PanFS file system (*para-sys); (ii*) Australia’s NCI national supercomputer facility, which uses a distributed Lustre file system (*dist-sys);* and (*iii*) an S3 bucket on Amazon AWS cloud (*cloud-sys*). Full computer specifications are provided in **Supplementary Table 2**.

When using BLOW5 and *Buttery-eel*, the basecalling rate (reads per second) scaled roughly linearly with the number of GPUs deployed (1-4 GPUs) on all three systems (**Figure 1b**), indicating that GPU capacity was the major constraint on overall performance. In contrast, when using FAST5 and *Guppy*, there was minimal improvement in the rate of basecalling with additional GPUs, indicative of a bottleneck in data access, rather than data processing (**Figure 1b**). As a result, we observed significant overall improvements in basecalling rates with BLOW5, compared to FAST5. The size of the improvement ranged from 2.1-fold (*cloud-sys*) to 5.3-fold (*dist-sys*) when using 4 GPUs (**Figure 1b**). Differences in the performance of the different systems reflect differences in their speed of FAST5 file reading; random access is slowest on distributed file systems, which therefore see greatest benefit from the switch to sequential access with BLOW5. Importantly, while the performance benefits of *BLOW5/Buttery-eel* varied depending on the system and the number of GPUs deployed, there was no scenario where it was outperformed by *FAST5/Guppy* (**Figure 1b**).

We next tested how these effects manifest during basecalling of a realistic human whole-genome sequencing dataset (at ~30X coverage; **Supplementary Table 1**). In FAST5 format, with 4 GPUs, this dataset took a minimum of 13.3 hours (*cloud-sys*) and a maximum of 41.6 hours (*dist-sys*) to be basecalled with *Guppy* (HAC model; **Figure 2c**). In BLOW5 format, basecalled with *Buttery-eel*, overall runtimes were reduced to ~5 hours on every system, corresponding to 2.7-fold (*cloud-sys*), 3.4-fold (*para-sys*) and 9.1-fold (*dist-sys*) improvements, respectively (**Figure 2c**). Notably, the differences in performance between systems during FAST5 basecalling, which is shaped by differences in the speed of random-access, is effectively erased by using BLOW5, because the analysis is now limited by the rate of data processing on the GPU, rather than file reading (**Figure 2c**).

Given that FAST5 files do not support sequential access, or multithreaded random access, an alternative path to improve the scalability of basecalling (or any other analysis) is to use multi–processing. ONT’s open-source development basecaller *Bonito* uses such a strategy. To evaluate multi-processing as a scalable approach to basecalling, we analysed the same small dataset as above in FAST5 format using *Bonito*. We also modified the *Bonito* source-code to permit sequential data access from BLOW5 files (see **Methods**). We then tested their performance on a local HPC environment with a network file system, using *Bonito’s* ‘fast-basecalling’ (FAST) model, chosen because *Bonito* is relatively slow compared to *Guppy*.

When basecalling a single dataset, we saw roughly equivalent run-times between *Bonito’s* multi-processing strategy with FAST5 and our sequential BLOW5 access strategy (1885 vs 1685 seconds; **Figure 2d,e**). Next, we emulated scenarios where the HPC’s file system is under increasing input/output (I/O) load by analysing multiple datasets simultaneously (n=1,2,4,8; see **Methods**). Whereas the time taken to basecall each individual dataset in BLOW5 format was unchanged regardless of how many datasets were processed in parallel, this was not true for FAST5 (**Figure 2d,e**). Instead, the rate of basecalling decreased and individual run-times for FAST5 datasets increased as the number of parallel runs (i.e. the I/O load) was increased (**Figure 2d,e**). When running eight datasets in parallel, each took ~5x longer to be processed with FAST5, compared to BLOW5, meaning a cumulative ~40x increase in overall execution time (**Figure 2e**). This result manifests from competition between random I/O operations, which exceed the capacity of the underlying hard disk drives (HDDs) during muti-processing. Therefore, while multi-processing can be used to improve performance in limited scenarios, it is not a viable strategy for high-throughput base-calling required for large genomics projects. In contrast, sequential data access on BLOW5 files is unaffected by computational load and therefore well suited to high-throughput basecalling.

## DISCUSSION

During large scale nanopore sequencing projects, basecalling is typically performed on external HPC or cloud computer environments, rather than the ONT sequencing device itself^3,5,7,8^. This is relatively slow, computationally expensive and is therefore a barrier to more widespread adoption of nanopore sequencing. Here, we show that the SLOW5 file format – developed previously to improve the efficiency of nanopore signal data analysis^9^ – can be used to significantly improve the performance of basecalling. *Buttery-eel* achieved 2.7–9.1-fold runtime improvements on a realistic human genome sequencing dataset processed on common HPC/cloud architectures, with identical outputs to the latest version of *Guppy*.

These performance gains are achieved by resolving a bottleneck in data access (I/O) caused by the FAST5 data format. However, data processing by the neural network basecalling model/s remains a costly operation that must be optimised to achieve further acceleration. This is especially true for ONT’s highest fidelity basecalling model, known as ‘super-accuracy’ (SUP), which is ~8 times slower than HAC in our experience, and therefore unfeasible for most users. ONT’s new basecalling software *Dorado* (currently in early prototype phase) is anticipated to introduce accelerated models in the future, and the availability of increasingly powerful GPUs will further accelerate data processing. In this context, it is important to note that the relative benefit of improved data access with BLOW5 will continue to increase as the efficiency of data processing improves. This is evident in a striking ~24-fold acceleration achieved by *Buttery-eel/BLOW5*, compared to *Guppy*/FAST5, when using ONT’s lightweight ‘fast’ basecalling model, where data processing is highly efficient. Further improvements could be made by direct integration of BLOW5 reading into future basecalling software, via the *slow5lib* library, rather than *Buttery-eel’s* indirect *Guppy* server approach.

*Buttery-eel* is the latest addition to the SLOW5 ecosystem, which already includes: (*i*) the SLOW5/BLOW5 file format and accompanying design specifications; (*ii*) the *slow5lib* (C/C++), *pyslow5* (python) and *slow5-rs* (rust) software libraries for reading and writing SLOW5/BLOW5 files; (*iii*) the *slow5tools* toolkit for creating, converting, handling and interacting with SLOW5/BLOW5 files^11^; and (*iv*) a suite of open source bioinformatics software packages with which SLOW5 is now integrated^12–17^. SLOW5 was conceived as an open-source, community-centric alternative to ONT’s FAST5 data format and the project continues to prioritize performance, compatibility, usability and transparency.

## METHODS & IMPLEMENTATION

### Reading SLOW5/BLOW5 files with *pyslow5*

The python library *pyslow5* is built on the *slow5lib* API using Cython which compiles the python code to C. It is designed to be easy to install, use, and to be fast. Though *pyslow5* is slower at reading and writing SLOW5/BLOW5 files than using C directly, it is still much faster than reading FAST5 files with the HDF5 python library, *h5py. Pyslow5* offers both random and sequential reading, as well as writing and appending of SLOW5/BLOW5 files. Each can be run with single threads, or with multiple threads, where the multithreading is controlled by function call flags and threading implemented on the C library side. Further multiprocessing from python can be applied to the random access methods, allowing for users to fine-tune resource usage.

### Design and implementation of *Buttery-eel*

*Buttery-eel* is open-source software written in python to enable basecalling of BLOW5 files directly with ONT’s production basecaller, *Guppy*. By using the software library *ont-pyguppy-client-lib* (https://pypi.org/project/ont-pyguppy-client-lib), *Buttery-eel* can control a *Guppy* server and *Guppy* client. Reads are read from a BLOW5 file and repackaged into a data structure compatible with the *ont-pyguppy-client-lib*, which can then be submitted to *Guppy* for basecalling. *Buttery-eel* thereby enables users to basecall BLOW5 files with the latest version of *Guppy*. The output sequence reads basecalled with *Buttery-eel* are identical to those basecalled with *Guppy. Buttery-eel* includes DNA methylation calling with built-in *Remora*, and writing output files in FASTQ or unaligned-SAM – both features of recent *Guppy* versions. Output files can also be split by mean phred quality score set by users into *pass* and *fail* files. A splitting tool *split_qscore.py* is also provided to allow users to split FASTQ or unaligned-SAM files on any given quality score.

*Buttery-eel* operates via the following internal workflow, which is summarized in **Figure 1a**:

1. Set up *Guppy* server;
2. Set up *Guppy* client to connect to *Guppy* server;
3. Read BLOW5 file with *pyslow5* library;
4. Submit reads to *Guppy* to be basecalled;
5. Get reads from *Guppy* and write to FASTQ/SAM.

### SLOW5 integration to Bonito

*Bonito* is an open-source research basecaller from ONT written in Python. It uses multiprocessing pools to extract reads from FAST5 files, assigns attributes from those reads to the Read class attributes, and passes that object to the basecalling module. Integrating SLOW5/BLOW5 access into *Bonito* was relatively straightforward using multithreaded sequential reading from the *pyslow5* library.

To integrate BLOW5 file usage with *Bonito*, three components were required:

1. Flags for setting SLOW5/BLOW5 format and controlling threads and batch size;
2. Assignment from *pyslow5* read object to Read class object;
3. Batching sequential reads to parse into multiprocessing pools for basecalling.

This integration effectively has C-level threads in the *pyslow5* library decompressing and reading reads sequentially, while using python multiprocessing to construct read objects and parse for basecalling. The FAST5 method however only has multiprocessing for both decompression, reading via random access, and constructing the read objects for basecall parsing.

### Basecalling performance benchmarks

The evaluation was performed on all systems, *dist-sys, para-sys*, and *cloud-sys* (**Supplementary Table 2**) for the small ~500,000 read dataset and the 30X whole genome dataset (**Supplementary Table 1**). Basecalling performance evaluations on FAST5 and BLOW5 were performed using ONT *Guppy* and on *Buttery-eel* (that wraps ONT *Guppy* server), respectively (see below). The small ~500,000 read dataset was basecalled with varying number of GPUs by setting `--device cuda:<devices>` option accordingly (`cuda:0` for one GPU, `cuda:0,1` for two GPUs, `cuda:0,1,2` for three GPUs and `cuda:0,1,2,3` for four GPUs). The whole genome dataset was basecalled using all four available GPUs. ONT *Guppy* for FAST5 input was executed with default options. *Buttery-eel* for BLOW5 was executed with the following additional parameters that are relevant to either BLOW5 or the server-client approach of *Guppy* (`ont-pyguppy-client-lib`).

- `--guppy_batchsize`: No. reads to send to the *Guppy* server at a time
- `--max_queued_reads`: Max capacity of *Guppy* server queue
- `--slow5_threads`: No. threads for decompressing and parsing BLOW5 file
- `--slow5_batchsize`: No. reads fetched from the BLOW5 file at a time
- `--procs`: No. worker processes to send/receive data from the *Guppy* server

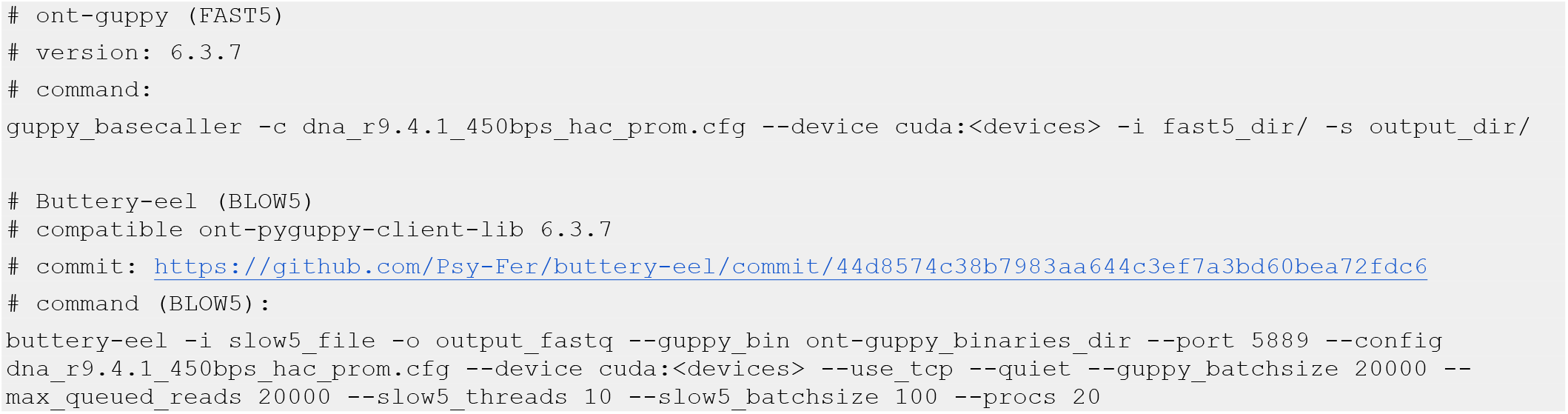

All experiments were performed using the high accuracy basecalling model. An additional experiment with the fast basecalling model was executed on *dist-sys* on the whole genome dataset using four GPUs. The execution times were measured using *GNU time* utility with *-v* options. The Linux disk cache (page cache, dentries, and inodes) was cleaned prior to each experiment, except on *dist-sys* due to lack of root access.

### Evaluating Bonito multi-processing

Bonito multi-processing approach was evaluated in an in-house local HPC system with a Network File System (NFS). The system consisted of a Synology DS3617xs Network Attached Storage (NAS) and two HPC server nodes. The NAS contained 12×12 TB hard disk drives configured under RAID 10 to form a EXT4 file system. The server nodes were connected to the NAS through 10 Gbps ethernet. Each server node has 4xTesla V100 GPUs. The NAS volume was mounted on both server nodes using NFS. Eight separate copies of the small ~500,000 read dataset (**Supplementary Table 1**) were made on the NAS. Then eight Bonito jobs were executed in parallel (4 jobs on each node, such that each job got exclusive access to a Tesla V100 GPU set using the environmental variable *CUDA_VISIBLE_DEVICES*), with each job accessing a separate copy of the data (out of the eight copies made). The aforementioned experiment was repeated for both FAST5 and BLOW5 inputs (see the commands below). The Linux disk cache (page cache, dentries, and inodes) were cleaned prior to each experiment on the NAS and servers. The execution time was measured using *GNU time* uitility with *-v* option. Bonito by default uses 8 multi-processes to fetch data from FAST5 files. For reading from BLOW5, `--slow5_threads` was set to 8 and `-- slow5_batchsize 4096` such that eight multi threads are used for decompressing and parsing a batch of 4096 BLOW5 records in parallel (sequence disk fetching is still done using a single thread).

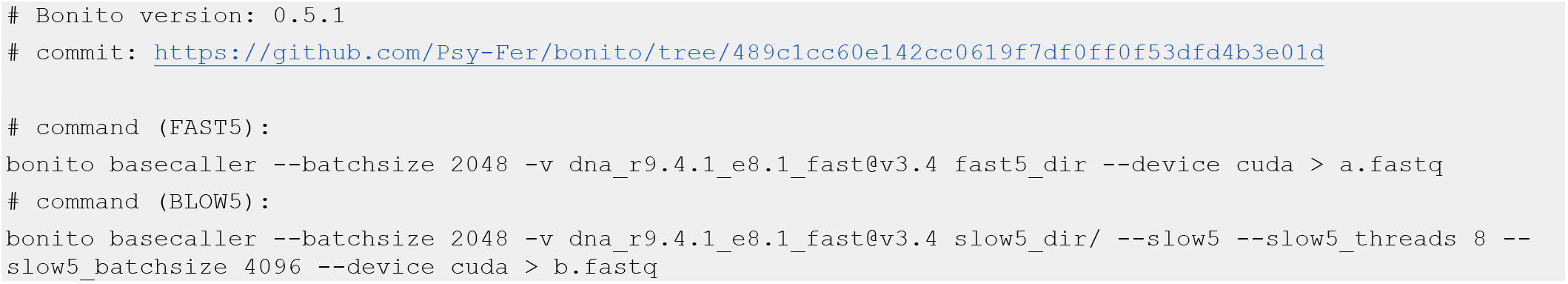

## DATA & CODE AVAILABILITY

Datasets used in benchmarking experiments are described in **Supplementary Table 1** and are available on NCBI Sequence Read Archive at Bioproject **PRJNA744329**. With the exception of ONT’s commercially available *Guppy* basecaller, all software used in this project is free and open source. SLOW5 and all of its associated software are free and open source:

Slow5tools: https://hasindu2008.github.io/slow5tools/

Slow5lib & pyslow5: https://hasindu2008.github.io/slow5lib/

Buttery-eel: https://github.com/Psy-Fer/buttery-eel

Bonito with SLOW5 support: https://github.com/Psy-Fer/bonito/tree/489c1cc60e142cc0619f7df0ff0f53dfd4b3e01d

SLOW5 format specification documents can be accessed at: https://hasindu2008.github.io/slow5specs

## ACKNOWLEDGEMENTS

We thank Alexander Payne for providing code examples and assistance for using the *Guppy* Python API. We thank Mark Bicknell from Oxford Nanopore Technologies for providing technical support with *ont-pyguppy-client-lib*. We thank Andre Luiz Martins Reis and Tim Amos for testing and providing early user feedback on *Buttery-eel*. We thank Derrick Lin for providing excellent technical support and freedom to use the institute’s HPC system in some quite exotic ways. Resources from the Australian National Computational Infrastructure were used during benchmarking experiments. We acknowledge the following funding support: Australian Medical Research Futures Fund grants MRF1173594 and MRF2016008 (to I.W.D.) and Australian Research Council DECRA Fellowship DE230100178 (to H.G.).

## DECLARATIONS

I.W.D. manages a fee-for-service sequencing facility at the Garvan Institute of Medical Research that is a customer of Oxford Nanopore Technologies but has no further financial relationship. H.G. & J.M.F. have previously received travel and accommodation expenses to speak at Oxford Nanopore Technologies conferences. H.G. and I.W.D. have paid consultant roles with Sequin PTY LTD. The authors declare no other competing financial or non-financial interests. The authors declare no other competing financial or non-financial interests.

## CONTRIBUTIONS

All authors contributed to the study design, figure generation and manuscript writing. H.S., J.M.F. and H.G. collaboratively developed *slow5lib, pyslow5, Buttery-eel*, modified the *Bonito* source code for BLOW5 integration, designed and executed benchmarking experiments, with supervision from I.W.D.

