## Supplementary Information for "Accelerated nanopore basecalling with SLOW5 data format"

### SUPPLEMENTARY TABLES

**Supplementary Table 1.** Datasets used for benchmarking experiments.

| Dataset description | Pore type | Reads | Total seq. (Gbases) | Total signal samples (Gsamples) | Median raw signal length per read (*n* signal samples) |
|---|---|---|---|---|---|
| PromethION human genome sample (NA12878, ~30X) | R9.4.1 | 9,083,052 | 93.4 | 1080.9 | 87,368 |
| Random subset of dataset **1** | R9.4.1 | 500,000 | 5.1 | 56.7 | 80,305 |

**Supplementary Table 2.** Computer specifications.

| System | Description | CPU (No. of cores/threads) | GPU type | RAM (GB) | File system | Disk system | OS |
|---|---|---|---|---|---|---|---|
| *dist-sys* | National Computer Infrastructure (NCI) GPU node | 2 x Intel Xeon Platinum 8268 (48/96) | 4 x Tesla V100-32 GB | 384 | Lustre | 7200 4TB disks in 120 NetApp disk arrays | CentOS 8.3.2011 |
| *para-sys* | Academic HPC with parallel file system | 2 x Intel Xeon Silver 4114 (20/40) | 4 x Tesla V100-16 GB | 384 | PanFS | ASH-100 12TB disks in RAID6+ configuration | CentOS 7.9.2009 |
| *cloud-sys* | AWS p3.8xlarge EC2 instance | 1 x Intel Xeon CPU E5-2686 v4 (16/32) | 4 x Tesla V100-16 GB | 240 | AWS S3 | - | Ubuntu 20.04.4 LTS |
